# Comparing Individual-Based Approaches to Modelling the Self-Organization of Multicellular Tissues

**DOI:** 10.1101/074351

**Authors:** James M. Osborne, Alexander G. Fletcher, Joseph M. Pitt-Francis, Philip K. Maini, David J. Gavaghan

## Abstract

The coordinated behaviour of populations of cells plays a central role in tissue growth and renewal. Cells react to their microenvironment by modulating processes such as movement, growth and proliferation, and signalling. Alongside experimental studies, computational models offer a useful means by which to investigate these processes. To this end a variety of cell-based modelling approaches have been developed, ranging from lattice-based cellular automata to lattice-free models that treat cells as point-like particles or extended shapes. It is difficult to accurately compare between different modelling approaches, since one cannot distinguish between differences in behaviour due to the underlying model assumptions and those due to differences in the numerical implementation of the model. Here, we exploit the availability of an implementation of five popular cell-based modelling approaches within a consistent computational framework, Chaste (http://www.cs.ox.ac.uk/chaste). This framework allows one to easily change constitutive assumptions within these models. In each case we provide full details of all technical aspects of our model implementations. We compare model implementations using four case studies, chosen to reflect the key cellular processes of proliferation, adhesion, and short-and long-range signalling. These case studies demonstrate the applicability of each model and provide a guide for model usage.

**Authors’ contributions:** JO and AF conceived of the study, designed the study, coordinated the study, carried out the computational modelling and drafted the manuscript. JP contributed to the computational modelling and helped draft the manuscript. PM and DG conceived of the study, designed the study and helped draft the manuscript. All authors gave final approval for publication.

## Introduction

Cells in eukaryotic organisms respond to physical and chemical cues through processes such as movement, growth, division, differentiation, death and secretion or surface presentation of signalling molecules. These processes must be tightly orchestrated to ensure correct tissue-level behaviour and their dysregulation lies at the heart of many diseases. The last decade has witnessed remarkable progress in molecular and live-imaging studies of the collective self-organization of cells in tissues. In combination with experimental studies, mathematical modelling is a useful tool with which to unravel the complex nonlinear interactions between processes at the subcellular, cellular and tissue scales from which organ- and organism-level function arises. The classical approach to modelling these processes treats the tissue as a continuum, using some form of homogenization argument to average over length scales much larger than the typical diameter of a cell. It can thus be difficult to incorporate heterogeneity between cells within a population, or investigate the effect of noise at various scales, within such models.

Facilitated by the reduction in cost of computing power, a number of discrete or ‘individual-based’ approaches have been developed to model the collective dynamics of multicellular tissues (Fig 1). Such models treat cells, or subcellular components, as discrete entities and provide natural candidates for studying the regulation of cell-level processes in tissue dynamics. However, they are less amenable to mathematical analysis than their continuum counterparts. The precise rules and methods of implementation differ between models and must be adapted to a particular biological system. However, they can be broadly categorised as on- and off-lattice, according to whether or not cells are constrained to lie on an artificial lattice. Each of the models described below have been helpful in furthering our knowledge but, like all models, they are simplifications and so have limitations.

**Fig 1.**
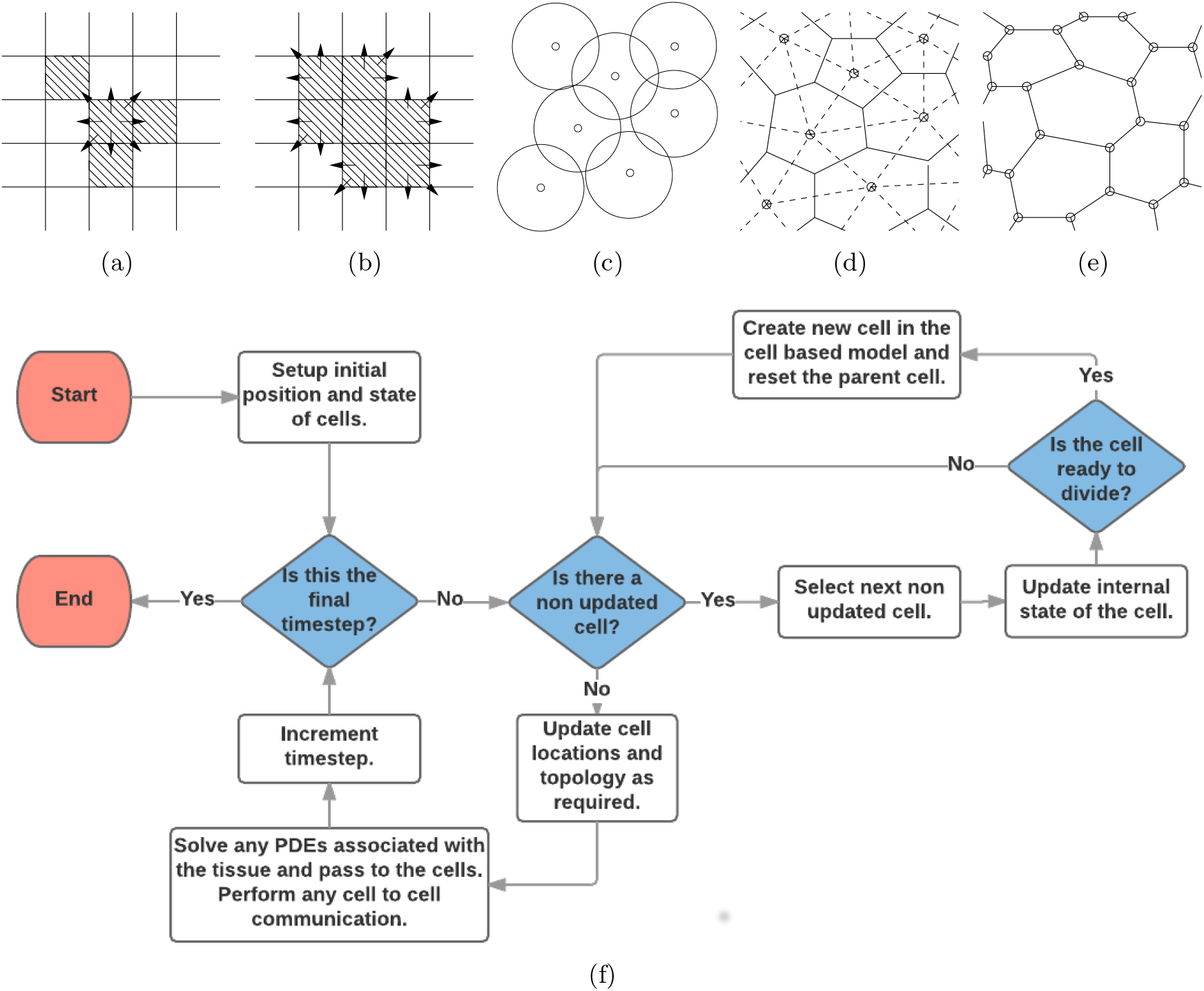
Schematics of the cell-based models considered in this study. (a) Cellular automaton (CA). (b) Cellular Potts (CP) model. (c) Overlapping spheres (OS) model. (d) Voronoi tessellation (VT) model. (e) Vertex model (VM). (f) Flow chart of cell-based simulation algorithm. See text for full details.

Arguably the simplest individual-based models are cellular automata (CA), where each lattice site can contain at most a single cell (Fig 1(a)). The system is evolved discretely, using a fixed time-stepping [1] or event-driven [2] approach, with the new state of each cell determined using deterministic or stochastic rules and the state of the system at the previous time step. The computational simplicity of CA renders them amenable to simulating large numbers of cells.

Another class of on-lattice model is the cellular Potts (CP) model [3], which represents each cell by several lattice sites, allowing for more realistic cell shapes (Fig 1(b)). The shape of each cell is evolved via some form of energy minimization. Unlike CA, the CP model can incorporate mechanical processes such as cell membrane tension, cell-cell and cell-substrate adhesion, chemotaxis and cell volume conservation. The CP model has been used to study biological processes ranging from cell sorting [4] and morphogenesis [5] to tumour growth [6].

The removal of a fixed-lattice geometry in off-lattice models enables the more detailed study of mechanical effects on cell populations. Two common descriptions of cell shape in off-lattice models are (i) ‘overlapping spheres’ (OS) or quasi-spherical particles [7] and (ii) through Voronoi tessellations (VT) [8]; in both approaches, the centre of each cell is tracked over time. In the former, cells are viewed as particles that are spherical in the absence of any interactions but which deform upon cell-cell or cell-substrate contact (Fig 1(c)). In the latter, the shape of each cell is defined to be the set of points in space that are nearer to the centre of the cell than the centres of any other cell; a Delaunay triangulation is performed to connect those cell centres that share a common face, thus determining the neighbours of each cell [9] (Fig 1(d)). In either case, Monte Carlo methods or Langevin equations may be used to simulate cell dynamics.

An alternative off-lattice approach commonly used to describe tightly packed epithelial cell sheets are vertex models (VM), in which each cell is modelled as a polygon, representing the cell’s membrane (Fig 1(e)). Each cell vertex, instead of centre, moves according to a balance of forces due to limited compressibility, cytoskeletal contractility and cell-cell adhesion. Additional rules govern cell neighbour rearrangements, growth, mitosis and death.

Several on- and off-lattice cell-based models have coupled descriptions of nutrient or morphogen transport and signalling to cell behaviour [10, 11, 12, 13, 14]. For example, a hybrid CA was used by Anderson and colleagues to study the role of the microenvironment on solid tumour growth and response to therapy [15], while Aegerter-Wilmsen et al. coupled a vertex model of cell proliferation and rearrangement with a differential algebraic equation model for a protein regulatory network to describe the interplay between mechanics and signalling in regulating tissue size in the *Drosophila* wing imaginal disk [10].

As the use of cell-based models becomes increasingly widespread in the scientific community, it becomes ever more useful to be able to compare competing models within a consistent computational framework, to avoid the potential danger of artifacts associated with different methods of numerical solution. To date there has not been a comparison of the classes of models described above, because it is difficult to identify in some cases corresponding processes but also that there has not been a common computational framework in which to carry out such a comparison. The development of Chaste, an open-source C++ library for cell-based and multiscale modelling [16, 17], now allows for the latter.

Here we present a systematic comparison of five classes of cell-based models through the use of four case studies. We demonstrate how the key cellular processes of proliferation, adhesion, and short- and long-range signalling can be implemented and compared within the competing modelling frameworks. Moreover, we provide a guide for which model is appropriate when representing a given system. We concentrate throughout on the two-dimensional case, but note that many of these models have also been implemented in three dimensions.

The remainder of this paper is structured as follows. We begin by presenting the five mathematical frameworks and discuss their implementation. Next, we use our four case studies to demonstrate how the modelling frameworks compare. Finally, we discuss our results and present a guide to which framework to use when modelling a particular problem.

## Materials and Methods

In this study we compare the implementation and behaviour of: cellular automata (CA); cellular Potts (CP); cell-centre, both overlapping sphere (OS) and Voronoi tessellation (VT); and vertex (VM) models. We begin by briefly describing the governing rules and equations for each of these models focussing on the way they implement the common processes of cell-cell interaction and cell division. Throughout, full references are given to previous publications giving much fuller details of the derivation and implementation of each of these models. We also present a consistent numerical implementation for the models.

### Cellular automaton (CA) model

There are several possible ways to represent cell movement in a CA. Here we focus on compact tissues so consider movement driven by division and cell exchange, using a shoving-based approach [18]. The spatial domain is discretised into a regular lattice with cells occupying the individual lattice sites (Fig 1(a)). The area *A*_*i*_ of each cell *i* in this model is given by 1 squared cell diameter (CD^2^).

In common with all of the cell-based models presented here, cell proliferation is determined by a model of how cells progress through the cell cycle, which in turn specifies when cells divide. Our model of cell-cycle progression varies across the four examples considered. However, in all cases a dividing cell selects a random lattice site from its Moore neighbourhood (the eight cells that surround it), and all cells along the row, column or diagonal from the dividing cell’s location are instantaneously displaced or ‘shoved’ to make space for the new cell.

We use a Metropolis-Hastings algorithm to make additional updates to the state of the tissue using asynchronous updating. At each time step ∆*t*, after checking for and implementing any cell divisions, we sample with replacement *N*_Cells_ cells, where *N*_Cells_ is the number of cells in the tissue at time *t* (thus, it may be the case that a cell is sampled more than once in a time step, while others are not sampled). This sweeping of the domain is also known as a Monte Carlo Step (MCS). We randomly select a neighbouring lattice site from each sampled cell’s Moore neighbourhood for a potential swap. The swapping of cells is intended to model random motility and the affinity of cells to form and break connections with adjacent cells. Assigning the MCS to a time step ∆*t* allows us to parametrize the timescale of the switching process and relate this to cell division. A probability per hour is assigned for the cells (or empty lattice site, which we refer to as a void) to swap locations, *p*_*swap*_, which is calculated as

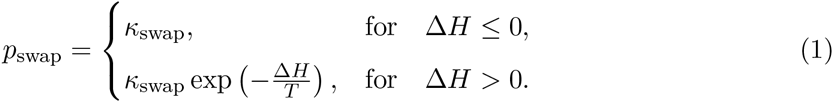

where *κ*_swap_ represents the rate of switching and *T* represents the background level of cell switching, modelling random cell fluctuations. If *T* = 0 then only energetically favourable swaps happen, and we use this as the default value for our simulations; as *T* increases, more energetically unfavourable swaps occur. Finally, ∆*H = H*_1_ − *H*_0_ denotes the change in adhesive energy due to the swap, with *H*_0_ and *H*_1_ being the energy in the original and changed configurations respectively, which is defined to be the sum of the adhesion energy between lattice sites:

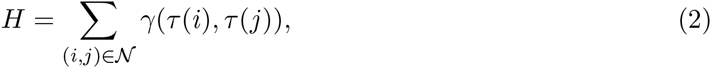

where *γ*(*a, b*) is a constant whose value depends on *a* and *b,* representing the adhesion energy between cells (or void) of type *a* and *b, τ*(*k*) is the type of cell *k* (or void if there is no cell on the lattice site) and 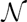 is the set of all neighbouring lattice sites. Here *τ*(*k*) takes the values ‘A’, ‘B’ and ‘void’, but can in principle be extended to more cell types.

### Cellular Potts (CP) model

As in the CA, we discretize the spatial domain into a lattice. Although, as in the CA case, the structure and connectivity of this lattice may be arbitrary, for simplicity we restrict our attention to a regular square lattice of size *N × N.* In contrast to the CA model, each cell is represented by a collection of lattice sites, with each site contained in at most one cell with the cell type of a lattice site being referred to as its spin. The area *A*_*i*_ of each cell *i* in this model is given by the sum of the area of all the lattice sites contained in the cell. In the present study, we take the area of each lattice site to be 1/16 CD^2^ and cells have a target area of 16 lattice sites i.e. 1 CD^2^. This is illustrated in Fig 1(b).

In a similar manner to the CA, the system evolves by attempting to minimize a total ‘energy’ or Hamiltonian, *H,* over discrete time steps using a Metropolis-Hastings algorithm. The precise form of *H* varies across applications but can include contributions such as cell-cell adhesion, hydrostatic pressure, chemotaxis and haptotaxis [5]. One iteration of the algorithm consists of selecting a lattice site and a neighbouring site (from the Moore neighbourhood) at random and calculating the change in total energy resulting from copying the spin of the first site to the second, ∆*H* = *H*_1_ − *H*_0_. The spin change is accepted with probability

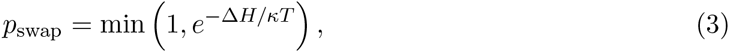

where *κ* is the Boltzmann constant and *T*, referred to as the ‘temperature’, characterizes fluctuations in the system; broadly speaking, at higher values of *T* cells move more freely, and hence system fluctuations increase in size. At each time step, ∆*t* we choose to sample with replacement *N × N* lattice sites (thus, it may be the case that a cell is sampled more than once in a time step, while others are not sampled). Note that this established algorithm for simulating CP models permits cell fragmentation, in principle; however, recent work has overcome this limitation [19].

In this study, we use a Hamiltonian of the form

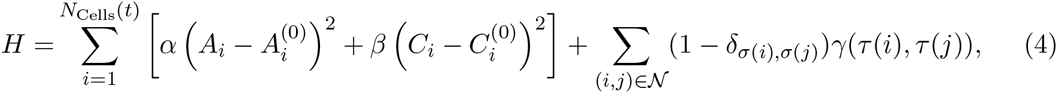

where the first and second terms on the right-hand side represent the area and perimeter constraint energies, summed over each cell in the system, and the third term represents the adhesion energy. Here *σ*(*k*) denotes the index of the cell containing lattice site *k* (note we let *σ*(*k*) = 0 if no cell is attached to the lattice site and we denote this to be the void), and *δ*_*a,b*_ is the delta function, which equals 1 if *a = b* and 0 otherwise. *τ*(*k*) denotes that cell’s ‘type’ (with the type void if *σ*(*k*) = 0), and *γ* denotes the interaction energies between cells occupying neighbouring lattice sites *i* and *j.* Again 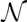 is the set of all neighbouring lattice sites and we allow *γ* to take different values for homotypic and heterotypic cell-cell interfaces and for ‘boundary’ interfaces between cells and the surrounding medium. Here 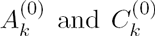 denote a specified target area and ‘target perimeter’ for cell *k*, respectively, which can depend on internal properties of the cell, allowing for cell growth to be modelled. Here we assume all cells are mechanically identical and set 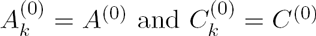. The parameters *α* and *β* influence how fast cells react to the area and perimeter constraints, respectively.

### Cell-centre models

Here cells are represented by their centres, which are modelled as a set of points 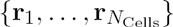 which are free to move in space. For simplicity, we assume all cells to have identical mechanical properties and use force balance to derive the equations of motion. We balance forces on each cell centre and making the standard assumption that inertial terms are small compared to dissipative terms (as cells move in dissipative environments of extremely small Reynolds number [20]), we obtain a first-order equation of motion for each cell centre, **r**_*i*_, given by

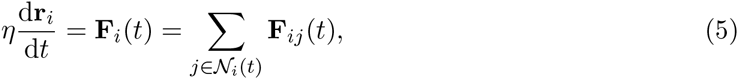

where *η* denotes a damping constant and **F**_*i*_(*t*) is the total force acting on a cell *i* at time *t* which is assumed to equal the sum of all forces, coming from the connections with all neighbouring cells 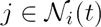 adjacent to *i* at that time, **F**_*ij*_(*t*). The definition of 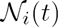 varies between the OS and VT models; in the former, it is the set of cells whose centres lie within a distance *r*_max_ from the centre of cell *i,* while in the latter, it is the set of cells whose centres share an edge with the centre of cell *i* in the Delaunay triangulation. We solve this equation numerically using a simple forward Euler scheme with sufficiently small time step ∆*t* to ensure numerical stability:

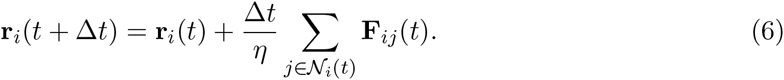

If the subcellular machinery causes cell *i* to divide then we generate a random mitotic unit vector 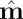 and the daughters cells are placed at 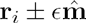, where ϵ is a constant division separation parameter and is dependent on the particular cell centre model being used.

### Overlapping spheres (OS)

Here, each cell *i* has an associated radius *R*_*i*_. Two cells *i* and *j* are assumed to be neighbours if their centres satisfy ||**r**_*i*_ - **r**_*j*_|| < *r*_max_ for a fixed constant *r*_max_, known as the interaction radius, where *r*_max_ > 2*R*_*i*_ for all *i.* The area of the cell is defined as [21]

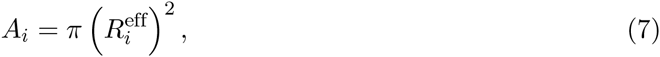

where

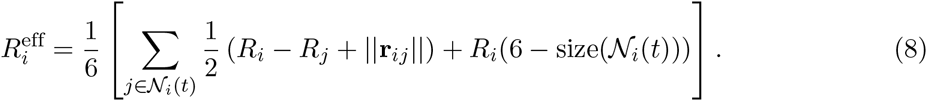

Here 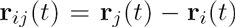 is the vector from cell *i* to cell *j* at time *t.* An illustration of cell connectivity is given in Fig 1(c).

In the OS model we define the force between cells as [21]

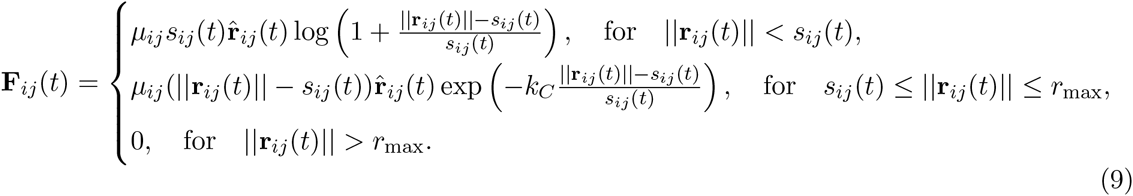

Here µ_*ij*_ is known as the “spring constant” and controls the size of the force (and depends on the cell types of the connected cells), by default *µ*_*ij*_ = *µ* for all interactions, 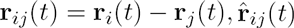, is the corresponding unit vector, *k*_*C*_ is a parameter which defines decay of the attractive force, and *s*_*ij*_(*t*) is the natural separation between these two cells. For the OS model *s*_*ij*_(*t*) is the sum of the radii of the two cells, and here the cell’s radius increases from 0.25 to 0.5 CDs over the first hour of the cell cycle, and hence is a function of time. Note that there is a cut off distance, *r*_max_, such that once 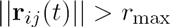 the cells are not connected so the force is zero.

### Voronoi tessellation (VT)

In the VT model we represent cells by the Voronoi region of the cell centres (this is defined as the region of space that is nearer to one cell centre than any other). Example cell regions are shown as solid lines in Fig 1(d). In this model, the area *A*_*i*_ of a cell *i* is defined to be the area of the corresponding Voronoi region. Cell connectivity is defined by the dual of the Voronoi region, known as a Delaunay triangulation and this is shown by the dashed lines in Fig 1(d). Two cell centres are assumed to be connected if they share an edge in the Delaunay triangulation.

In the VT model we define the force between cells to be [8, 22],

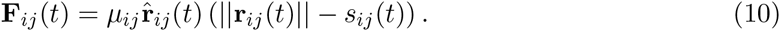

Here *µ*_*ij*_ is the spring constant (which again defaults to *μ*_*ij*_ = *μ*), 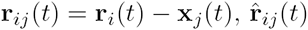 is the corresponding unit vector and *s*_*ij*_(*t*) is the natural separation between these two cells. For the VT model this increases linearly from 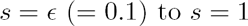 over the first hour of the cell cycle.

When using a Delaunay triangulation to define cell connectivity, on a growing tissue, long edges can form between distant cells causing unrealistic connections to be made. There are two methods used to overcome this. The first is to introduce a cut-off length, *r*_max_, such that cells further apart than the cut-off length are no longer connected (analogous to the OS model). The second method is to introduce *ghost nodes,* which are extra nodes introduced into the simulation which surround the tissue, which do not exert any forces on the cells, and preclude any long connections from forming. Moreover these ghost nodes ensure that the Voronoi regions, and therefore cell areas, are finite. In order for the ghost nodes to surround the tissue, as it grows, cells exert a force on the ghost nodes (and ghost nodes exert forces on other ghost nodes) causing them to move with the cells. The force applied is calculated using Equation (10). For more details on ghost nodes see [23].

### Vertex model (VM)

In the VM a tissue is represented by a collection of non-overlapping connected polygons whose vertices are free to move, each polygon corresponds to a cell. In this model, the area *A*_*i*_ of a cell *i* is given by the area of the associated polygon. An illustration of cells in a VM is given in Fig 1(e). As in cell-centre models we consider a set of points 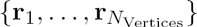 and we use force balance to derive an equation of motion [24]:

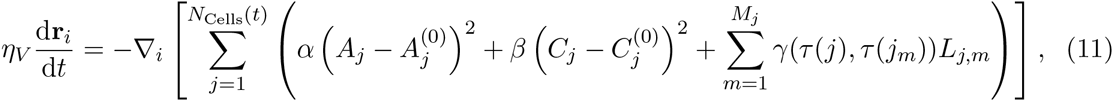

where **r**_*i*_ is the position of vertex 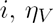 is an associated drag constant, 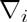 is the gradient with respect to **r**_*i*_ and *N*_Cells_(*t*) denotes the number of cells in the system at time *t*. The variables *A*_*j*_ and *C*_*j*_ denote the cross-sectional area and the perimeter of cell *j,* respectively, and *M*_*j*_ is the number of vertices of cell *j*. *L*_*j,m*_ is the length of the line connecting vertices *m* and *m* + 1 in cell *j* and *j*_*m*_ is the neighbour of cell *j* which shares the edge connecting vertices *m* and *m* + 1 in cell *j.* Similar to the CP model, *A*^(0)^ is the cell’s natural (or target) area, and *C*^(0)^ is its natural perimeter. Finally, *α* and *β* are positive constants that represent a cell’s resistance to changes in area or perimeter, respectively. *γ* again denotes the interaction energies between neighbouring cells. We allow *γ* to take different values for homotypic and heterotypic cell-cell interfaces and for ‘boundary’ interfaces between cells and the surrounding medium.

For simplicity here we set all cells to have a target area of *A*^(0)^ = 1 and therefore a target perimeter of 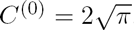. See [25] for a discussion on the other growth options and their influence on simulations.

To maintain a non-overlapping tessellation of the domain we need to introduce a process where cell edges can swap, known as a T1 transition. This process allows cell connectivity to change as cells grow and move and is instrumental in the process of cell sorting. When an edge between two cells, *A* and *B,* becomes shorter than a given threshold, *l*_*r*_, we rearrange the connectivity so that the cells *A* and *B* are no longer connected and the other cells that contain the vertices on the short edge, *C* and *D,* become connected. Other processes may also be required, such as a T2 transition where small triangular elements are removed to simulate cell death. For further details of these elementary operations, see [26].

As with all of these models, other force laws could be used to define cell interactions [27]. For full details of the forces used in the vertex model, along with how they differ in both implementation and simulation results, see [26].

### Implementation

Now we have briefly introduced all the cell-based models used in this study we proceed to discuss their implementation. Each simulation takes the form given in Fig 1(f). All components of this algorithm are the same for each simulation type except for the CA model where cells may also move due to the division of other cells. All models have been non-dimensionalised so that the units of space are cell diameters (CDs) and time is measured in hours.

We implement all model simulations within Chaste, an open source C++ library that provides a systematic framework for multiscale multicellular simulations [17]. Further details on the implementation of VM and CP models within Chaste can be found in [26] and [28], respectively.

## Results

We now present a series of case studies that illustrate how cellular processes can be represented in each cell-based model and how differences in representation may influence simulation results.

### Adhesion

Cell-cell adhesion is a fundamental property of tissue self-organization. If embryonic cells of two or more histological types are placed into contact with each other, they can undergo spontaneous reproducible patterns of rearrangement and sorting. This process can, for example, lead to engulfment of one cell type by another. Explanations for this phenomenon include the differential adhesion hypothesis, which states that cells tend to prefer contact with some cell types more than others due to type-specific differential intercellular adhesion [29]; and the differential surface contraction hypothesis, which states that cells of different types instead exert different degrees of surface contraction when in contact with other cell types or any surrounding medium [30]. Computational modelling has played a key role in comparing these hypotheses [31].

As our first case study, we simulate cell sorting due to differential adhesion in a monolayer of cells in the absence of cell proliferation or respecification. We consider a mixed population of two cell types, A and B, which we assume to exhibit differential adhesion. This is implemented in the CA, CP and VM models by having different values of the parameter *γ* for different cell types. Specifically, we choose *γ*(A, A) = *γ*(B, B) < *γ*(A, B) and *γ*(A, void) < *γ*(B, void) to drive type-A cells to engulf type-B cells. In the cell-centre (OS and VT) models, we instead assume that for any pair of neighbouring cells located a distance farther apart than the rest length, the spring constant, *µ* is reduced by a factor *µ*_*het*_ = 0.1 if the cells are of different types. Additionally, in the OS model we use a larger interaction radius, *r*_max_ = 2.5, to encourage cell sorting.

In addition to the update rules and equations of motion outlined in the previous section, we consider each cell to be subject to random motion. This random motion is intrinsic to the CA and CP models and is adjusted by changing the parameter *T* in Eq (1) and Eq (3). For the OS, VT and VM models we introduce an additional random ‘diffusive’ force acting on each cell or vertex,

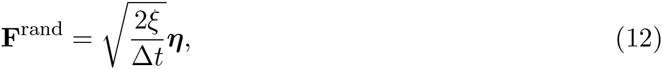

where *η* is a vector of samples from a standard multivariate normal distribution and *ξ* is a parameter that represents the magnitude of the perturbation [26]. This size is scaled by the time step to ensure that when the equations of motion are solved numerically, the rate of diffusion is independent of the size of time step. We simulate each model ten times, starting from an initial rectangular domain of width *L*_*x*_ and height *L*_*y*_, comprising 50% type-A cells and 50% type-B cells. For all models, the edge of the domain is a free boundary.

The time step of the CA and CP models dictates how many MCS occur per hour and, along with the temperature, *T*, can influence the dynamics of the simulation [28]. Here we perform an *ad hoc* calibration of *T* and ∆*t* so that the temporal dynamics of the CA and CP models match those of the other models as far as possible [28]. A full list of parameter values is provided in Tables 1 and 2.

**Table 1.** Table of parameters used in the models across case studies.

| Parameter | Description | Model(s) | Value | Reference |
| --- | --- | --- | --- | --- |
| $\Delta t$ | Time step | CA, CP | 0.01 h | [28]* |
|  |  | OS, VT, VM | 0.005 h | [26][22] |
| $A^{(0)}$ | Cell target area | CA | 1 ls (1.0 CD<sup>2</sup>) | – |
| | | OS | $\pi/4$ CD<sup>2</sup> | – |
|  |  | CP | 16 ls (1.0 CD<sup>2</sup>) | – |
| | | VT | $\sqrt{3}/2$ CD<sup>2</sup> | – |
|  |  | VM | 1 CD<sup>2</sup> | [26] |
| $\gamma(\text{cell, cell})$ | Cell-cell adhesion coefficient | CA, CP | 0.1 | [28]* |
|  |  | VM | 1 | [26] |
| $\gamma(\text{cell, void})$ | Cell-boundary adhesion coefficient | CA, CP | 0.2 | [28]* |
|  |  | VM | 10 | [26] |
| $k_C$ | Decay of attraction force | OS | 5 | [21] |
| $T$ | Temperature | CA | 0.0 | – |
|  |  | CP | 0.1 | [28] |
| $\alpha$ | Volume deformation coefficient | CP | 0.1 | [28] |
|  |  | VM | 50.0 | [26] |
| $\beta$ | Surface deformation coefficient | CP | 0.01 | [28] |
|  |  | VM | 1.0 | [26] |
| $C^{(0)}$ | Cell target perimeter | CP | 16 ls (4 CD) | [28] |
| | | VM | $2\sqrt{\pi}$ CD | [26] |
| $\eta$ | Drag coefficient | OS, VT | 1.0 | [22] |
|  |  | VM | 1 | [26] |
| $\mu$ | Spring constant | OS, VT | 50.0 | [8, 32] |
| $s$ | Mature cell spring rest length | OS, VT | 1.0 CD | [22] |
| $\kappa_{\text{swap}}$ | rate of cell switching | CA | 1 | – |
| $r_{\text{max}}$ | Force cut-off length | OS | 1.5 | – |
| $l_r$ | Cell rearrangement threshold | VM | 0.1 CD | [26] |

Asterisks (\*) denote parameters whose values are taken for the CP model from the given reference, with the same value being assumed for the CA model.

**Table 2.** Table of parameters specific to the differential adhesion simulations.

| Parameter | Description | Model(s) | Value | Reference |
| --- | --- | --- | --- | --- |
| $L_x$ | Initial width of tissue | All | 10 CD | — |
| $L_y$ | Initial height of tissue | All | 10 CD | — |
| $t_{\text{cycle}}$ | Mean cell cycle duration | All | 16 h | — |
| $t_{\text{end}}$ | Simulation duration | All | 100 h | — |
| $\gamma(\text{A}, \text{B})$ | Heterotypic cell-cell adhesion coefficient | CA | 0.2 | * |
|  |  | CP | 0.5 | * |
|  |  | VM | 2 | * |
| $\gamma(\text{B}, \text{void})$ | Type B cell-boundary adhesion coefficient | CA | 0.4 | * |
|  |  | CP | 1.0 | * |
|  |  | VM | 20 | * |
| $T$ | Base ‘temperature’ | CA | 0.1 | * |
|  |  | CP | 0.2 | [28]* |
| $\mu_{\text{het}}$ | Heterotypic spring constant | OS, VT | 0.1 | [33] |
| $r_{\text{max}}$ | Force cut-off length | OS | 2.5 | * |
| $\xi$ | Level of noise | OS | 0.05 | * |
|  |  | VT | 0.1 | * |
|  |  | VM | 0.1 | * |

Asterisks (\*) denote parameters whose values in the CA and CP models are chosen to ensure that engulfment occurs over a similar timescale to that observed in the OS, VT and VM models.

The results of a single simulation of each model are shown in Fig 2. In each case, the tissue evolves to a steady state where cells of each type are more clustered than the initial configuration. In the CA, CP and VM models, type-A cells are eventually completely engulfed; note that for other parameter values, each model can exhibit dissociation or checkerboard patterning [4, 31]. In the other models, the tissue evolves to a local steady state that does not correspond to complete engulfment.

**Fig 2.**
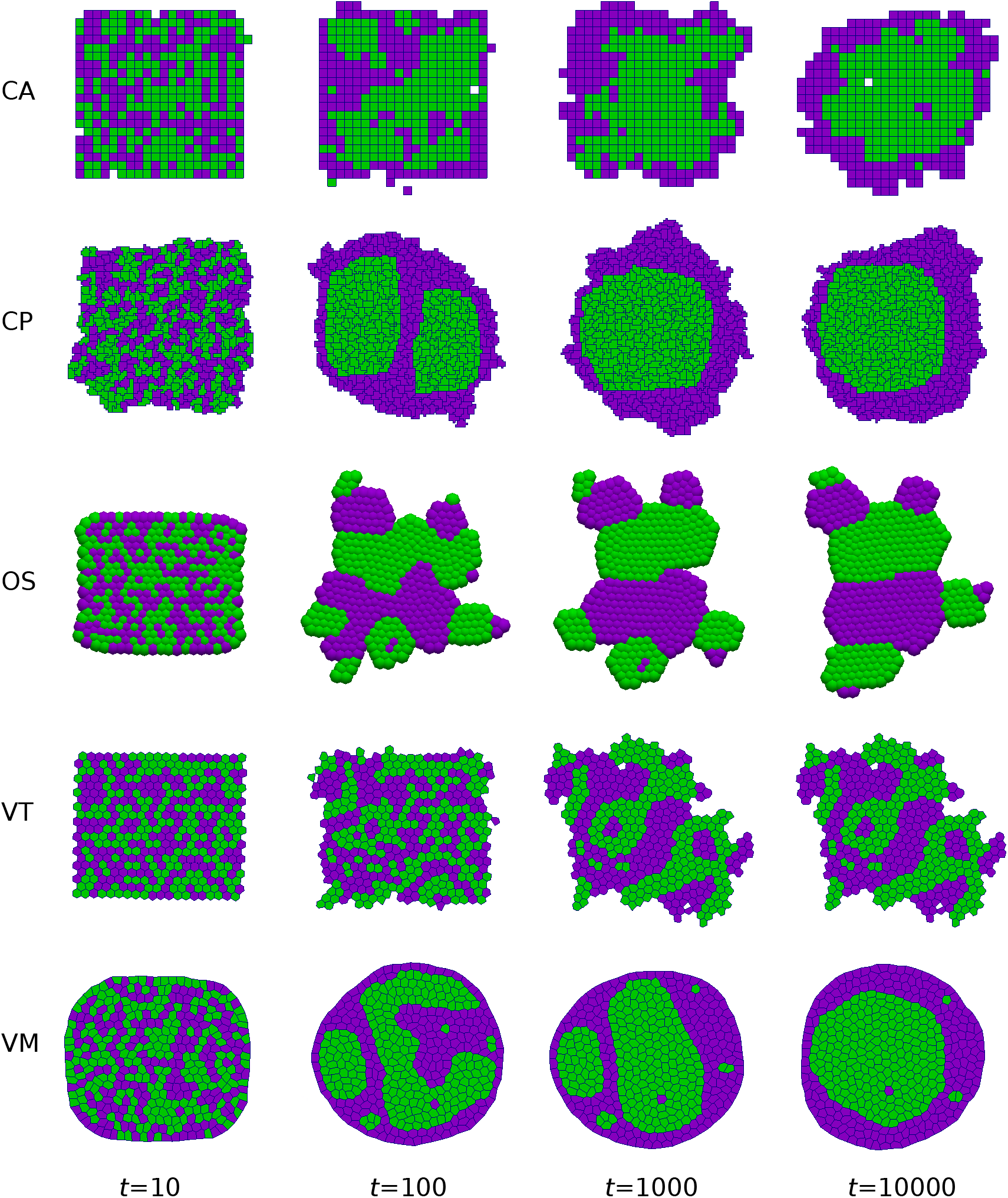
Simulations of cell sorting due to differential adhesion. Snapshots are shown at selected times for each model. Cells of type A and B are shown in purple and green, respectively. Engulfment of type-B cells occurs most readily in the CA, CP and VM models. Parameter values are given in Tables 1 and 2.

A quantitative comparison of cell sorting dynamics is shown in Fig 3. In particular we show how cell sorting is affected by the level of random motion applied to cells. This is demonstrated by computing the fractional length, defined as the total length of edges between cells of different types for each simulation. These are then normalised by the length at *t* = 0 for comparison. We find that the CA and CP models undergo repeated annealing due to their stochastic updating, and eventually end up at the global minimum (corresponding to complete engulfment). However, large amounts of noise can cause disassociation of cells in the CP model.

**Fig 3.**
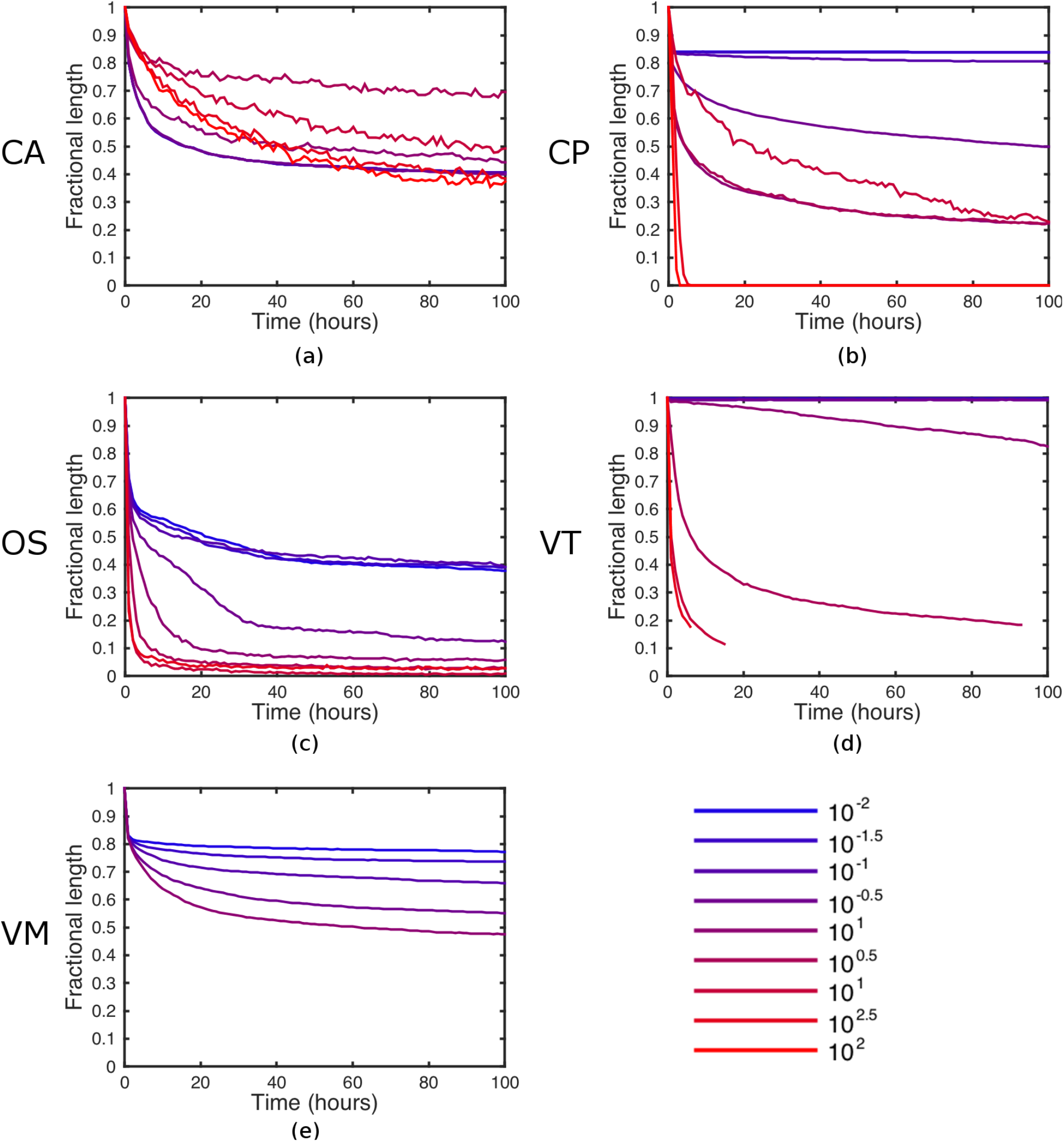
Comparison of cell sorting dynamics across differential adhesion simulations. As a measure of sorting, the fractional length is computed as a function of time for each model: (a) CA; (b) CP; (c) OS; (d) VT; (e) VM. Results are shown for varying multiples of the baseline level of noise, *ξ*, whose value is defined for each model in Table 2. Each line is the mean value of 10 simulations. Parameter values are given in Tables 1 and 2.

As Fig 3 shows, for the off-lattice models the total energy of the system evolves to a local minimum in the absence of noise; however, we can recover more complete engulfment through the addition of random cell movement. A relatively large amount of noise is required to alter cell neighbours in the Delaunay triangulation, illustrated by the flat lines in Fig 3(b). However, if there is too much noise then cells can become dissociated and move amongst the ghost nodes; in this case, if a cell reaches the edge of the ghost node region, its Voronoi area becomes ill-defined. A similar sensitivity is exhibited by the VM; in this case, if the amount of noise is too high, cell shapes can become inverted due to vertices randomly intersecting edges.

To summarise, we find that the degree of cell sorting observed in our simulations depends on how much random cell movement can be accommodated within each model. We note that there is no reason *a priori* to suppose that the configuration corresponding to the global minimum is biologically realistic; this depends on how the typical time scale over which complete sorting occurs compares to other embryogenic processes.

### Proliferation, death and differentiation

Embryonic development and adult tissue self-renewal both rely on careful control of cell proliferation, differentiation and apoptosis to ensure correct cell numbers. The intestinal epithelium offers a particularly important example of such tightly orchestrated cell dynamics. It is folded to form invaginations called crypts and (in the small intestine) protrusions called villi. The disruption of cell proliferation and migration in intestinal crypts is the cause of colorectal cancers. Experimental evidence indicates a complex pattern of cell proliferation within the crypt, in which cells located at the base of the crypt cycle significantly more slowly than those further up. One possible explanation for this is contact inhibition, in which stress due to overcrowding causes a cell to proliferate more slowly, enter quiescence or even undergo apoptosis [34]. The biological mechanism through which shear stress affects the expression of key components in the Wnt signalling pathway, which in turn plays an important role in cell proliferation and adhesion in this tissue, has been elucidated through a number of studies [35, 36].

A variety of cell-based models have been developed to study aspects of intestinal crypt dynamics [37], including defining the role of the Wnt signalling pathway [38]. The process and consequences of contact inhibition have also been described using cell-based modelling approaches in a more general setting [39, 40, 41]. A recent study used a cell-centre modelling approach to investigate how combined changes in Wnt signalling response and contact inhibition may induce altered proliferation in radiation-treated intestinal crypts [33].

As our second case study, we simulate the spatiotemporal dynamics of clones of cells within a single intestinal crypt. This example demonstrates how multicellular models and simulations (in particular Chaste) can include the coupling of cell-level processes to simple subcellular processes and deals with cell proliferation, death and differentiation.

Our underlying model of a colonic crypt has been described in detail previously [23, 42, 43]. We restrict cells to lie on a fixed cylindrical crypt surface, defined by the two-dimensional domain [0, *L*_*x*_] × [0, *L*_*y*_], where *L*_*x*_ and *L*_*y*_ denote the crypt’s circumference and height, respectively. Periodicity is imposed at the left- and right-hand boundaries *x* ∈ {0, *L*_*x*_}. We impose a no-flux boundary condition at the crypt base (*y* = 0) and remove any cell that reaches the crypt orifice (*y* = *L*_*y*_). In each simulation, we start with a random tessellation of cells occupying this domain; the crypt is then evolved for a duration *t*_*start*_ to a dynamic equilibrium, before cell clones are recorded and the crypt evolved for a further duration *t*_*end*_.

For each cell-based model considered, we implement cell proliferation and differentiation as follows. Any cell located above a threshold height *y*_prolif_ from the crypt base is considered to be terminally differentiated, and can no longer divide. Any cell located below *y*_prolif_ can proliferate. On division a random cell cycle duration is drawn independently for each daughter cell. Specifically, we draw the duration of each cell’s G1 phase, *t*_*G*1_, from a truncated normal distribution with mean *µ*_G1_ = 2, variance 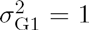 and lower bound 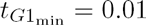, and we set the remainder of the cell cycle as *t*_*S*_ = 5, *t*_*G*2_ = 4 and *t*_*M*_ = 1, for the durations of the S phase, G2 phase, and M phase, respectively.

In addition the duration of G1 phase depends on the local stress, interpreted as the deviation from a cell’s preferred area. A cell pauses in the G1 phase of the cell cycle if

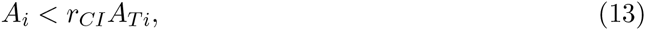

where *r*_*CI*_ is the quiescent area fraction and *A*_*i*_ is as earlier defined for each model [44]. This description allows for quiescence imposed by transient periods of high compression, followed by relaxation. If a cell is compressed during the G2 or S phases then it will still divide, and thus cells whose areas are smaller than the given threshold may still divide.

The dimensions of the crypt domain are chosen in line with [32] but are scaled to decrease simulation time. A full list of parameter values is provided in Tables 1 and 3.

**Table 3.** Table of parameters specific to the colonic crypt simulations.

| Parameter | Description | Model(s) | Value | Reference |
| --- | --- | --- | --- | --- |
| $t_{start}$ | Pre-simulation duration | All | 100 h | — |
| $t_{end}$ | Simulation duration | All | 1000 h | — |
| $r_{CI}$ | Quiescent volume fraction | All | 0 – 1 | — |
| $L_x$ | Width of crypt | All | 15 CD | Scaled from [32] |
| $L_y$ | Height of crypt | All | 12 CD | Scaled from [32] |
| $h$ | Height at which cells are sloughed | All | 12 CD | Scaled from [32] |
| $\mu_{G1}$ | G1 phase duration mean | All | 2 h | — |
| $\sigma_{G1}^2$ | Cell cycle variance | All | 1 h | — |
| $t_{G1_{min}}$ | Minimum G1 Phase duration | All | 0.01 h | — |
| $t_S$ | S Phase duration | All | 5 h | — |
| $t_{G2}$ | G2 Phase duration | All | 4 h | — |
| $t_M$ | M Phase duration | All | 1 h | — |
| $y_{prolif}$ | Proliferation height threshold | All | 6 CD | Scaled from [32] |

The results of a single simulation of each model are shown in Fig 4. In each case, the number of clones decreases over time as the crypt drifts to monoclonality. A more quantitative comparison of clonal population dynamics is shown in Fig 5. For each simulation we compute the number of clones remaining in the crypt as a function of time. All models exhibit the same qualitative behaviour, with a sharp initial drop as all clones corresponding to cells outside the niche are rapidly lost, followed by a more gradual decay in the number of clones at the crypt base due to neutral drift. However, we note that the number of clones reduces more slowly in the VM than other models, since the implementation of the ‘no flux’ boundary condition at the crypt base causes cells to remain there for longer in this model. This highlights the effect that the precise implementation of boundary conditions can have in such models. Finally, we note that for models where contact inhibition can be imposed, we see a slight effect of the degree of contact inhibition on the clonal population dynamics. In most of the models contact inhibition slows the process of monoclonal conversion, due to there being more compression at the crypt base. In contrast, in the VM the number of clones present in the crypt decreases more quickly when *r*_*CI*_ is larger. This effect is due to there being higher rates of division, resulting in cells more frequently being ‘knocked’ from the crypt base; in the other models this effect is counteracted by compression from above.

**Fig 4.**
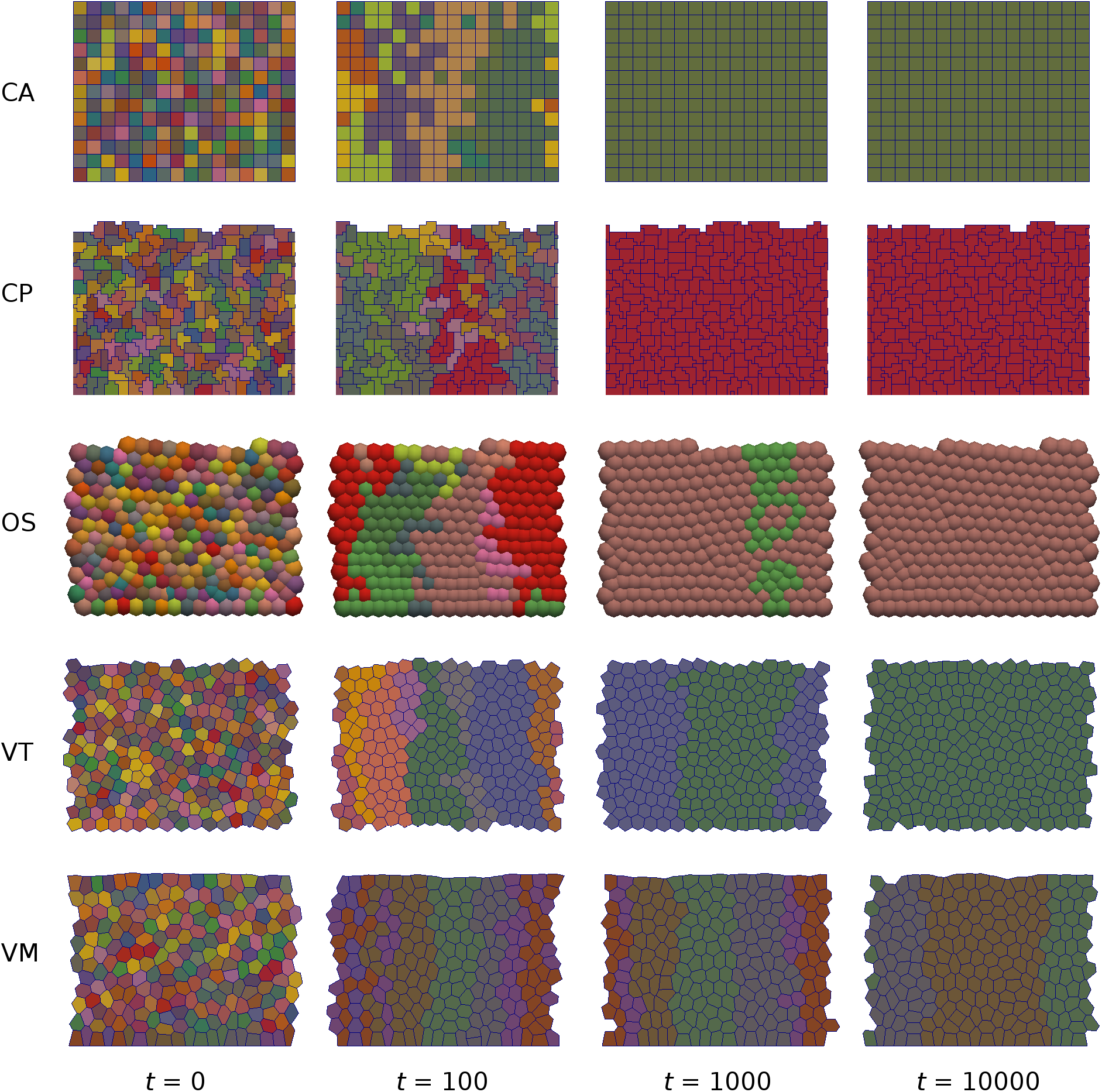
Simulations of monoclonal conversion in the colonic crypt. Snapshots are shown at selected times for each model. In each simulation at time *t* = 0, every cell is regarded as a clonal population and given a different colour, which is inherited by its progeny. These populations evolve in time due to cell proliferation and sloughing from the crypt orifice, resulting in a single clone eventually taking over the entire crypt. Parameter values are given in Tables 1 and 3.

**Fig 5.**
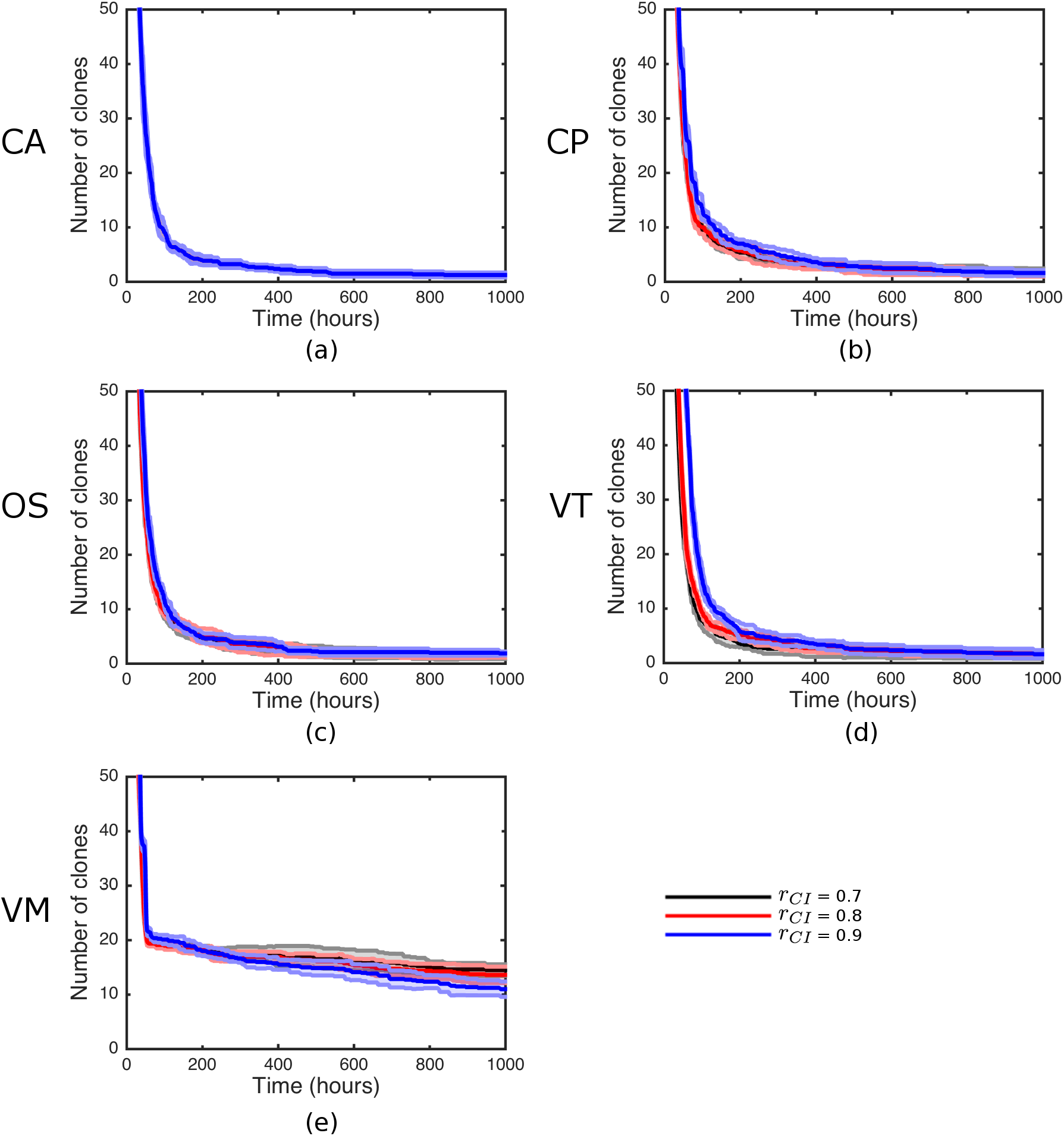
Comparison of clonal population dynamics across crypt simulations for varying levels of contact inhibition. The number of clones remaining in the crypt is computed as a function of time for each model: (a) CA; (b) CP; (c) OS; (d) VT; (e) VM. For each model, the mean and standard error from 10 simulations are shown for three levels of contact inhibition, quantified by the parameter *r*_*CI*_. Parameter values are given in Tables 1 and 3.

A quantitative comparison of cell velocity profiles up the crypt is shown in Fig 6. This extends the comparison previously made of cell-centre and vertex models of crypt dynamics in [42]. For each simulation we compute the vertical component of cell velocity at different heights up the crypt, averaging over the x direction. We find that all models are similar when considering a ‘position-based’ cell-cycle model (in which cell proliferation occurs below a threshold height up the crypt, corresponding to a threshold Wnt stimulus). However we see more pronounced differences when incorporating more restrictive contact inhibition into the cell-cycle model.

**Fig 6.**
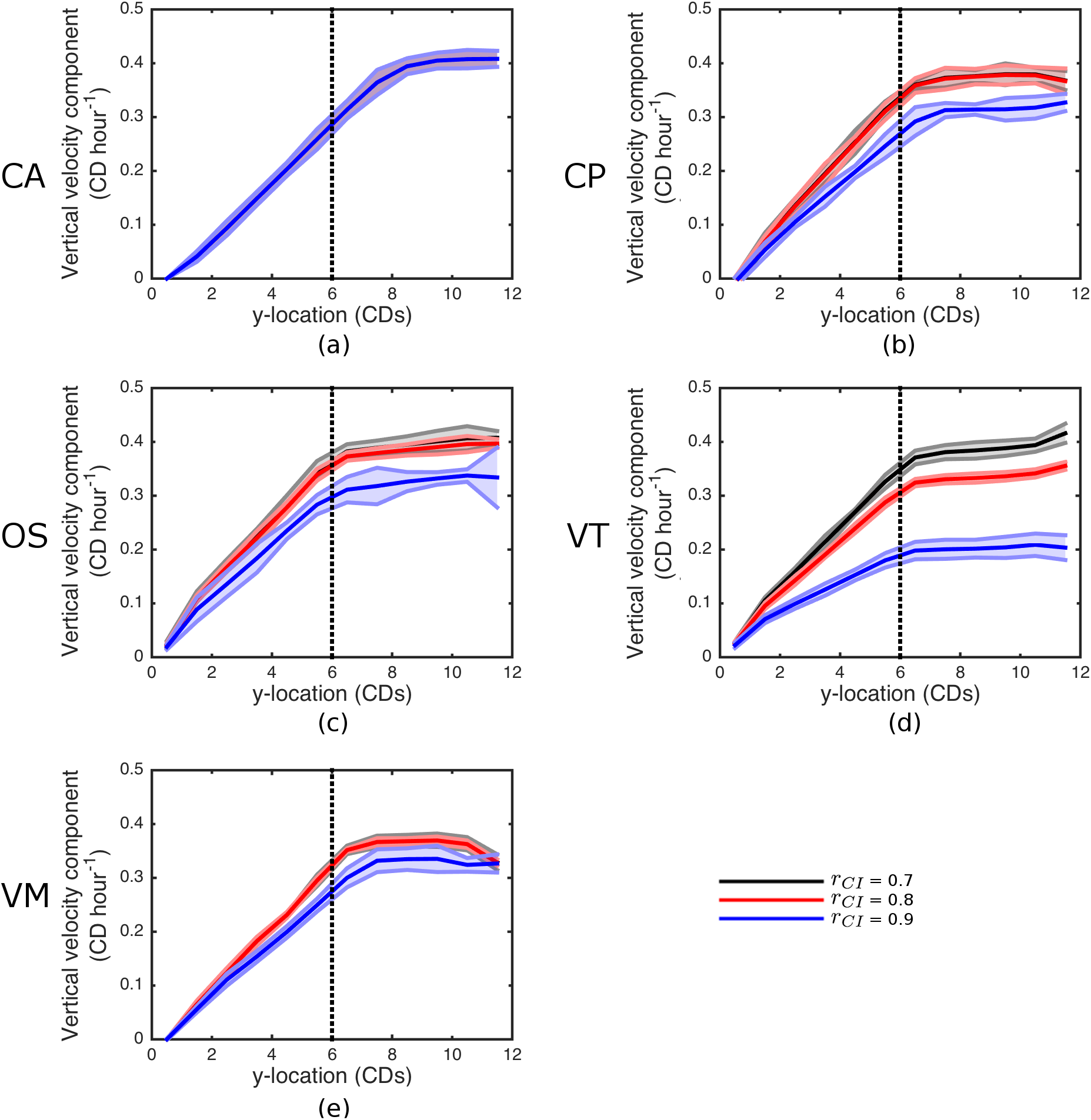
Comparison of crypt cell velocity profiles across simulations. The vertical component of cell velocity is computed for each model: (a) CA; (b) CP; (c) OS; (d) VT; (e) VM. For each model, the mean and standard error from 10 simulations are shown for three levels of contact inhibition, quantified by the parameter *r*_*CI*_. Parameter values are given in Tables 1 and 3.

### Short-range signalling

In many developmental processes, distinct states of differentiation emerge from an initially uniform tissue. Lateral inhibition, a process whereby cells evolving towards a particular fate inhibit their immediate neighbours from doing so, has been proposed as a mechanism for generating such patterns. This process is known to be mediated by the highly conserved Notch signalling pathway, which involves ligand-receptor interactions between the transmembrane proteins Notch and Delta or their homologues [45].

Lateral inhibition through Notch signalling has been the subject of several mathematical modelling studies [46, 47, 48, 49, 50, 51]. Such models have largely focused on the conditions for fine-grained patterns to occur in a fixed cell population; little attention has been paid to its interplay with cell movement, intercalation and proliferation. To illustrate how cell-based modelling approaches may be utilised to investigate such questions, as our third case study we simulate Notch signalling in a growing monolayer, with cell proliferation dependent on Delta levels. This example demonstrates how intercellular signalling may be incorporated within each cell-based model.

In this example, cells proliferate if located within a radius *R*_*P*_ from the origin, and are removed from the simulation if located more than a radius *R*_*S*_ > *R*_*P*_ from the origin. For each proliferative cell, we allocate a probability *p*_div_ of division per hour, once the cell is above a minimum age, *t*_min_. This is implemented by independently drawing a uniform random number *r* ~ *U*[0,1] for each cell at each time step and executing cell division if *r* < *p*_div_∆*t*.

This description is coupled to a description of Notch signalling between neighbouring cells that is based on a simple ordinary differential equation model previously developed by Collier et al. [46]. This represents the temporal dynamics of the concentration of Notch ligand, *N*_*i*_(*t*), and Delta receptor, *D*_*i*_(*t*), in each cell *i* in the tissue. A feedback loop is assumed to occur, whereby activation of Notch inhibits the production of active Delta. Signalling between cells is reflected in the dependence of Notch activation on the average level of Delta among a cell’s immediate neighbours. The precise set of equations for this signalling model takes the form

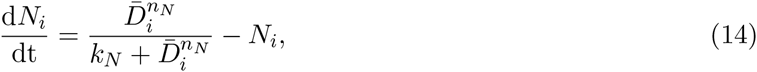

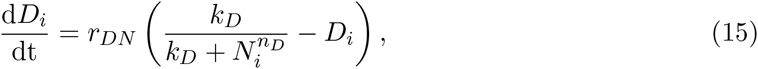

where 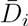 denotes the average value of 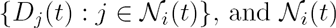 is the set of neighbours of cell *i.* A full list of parameter values is provided in Tables 1 and 4. At the start of the simulation, values of each *N*_*i*_ and *D*_*i*_ are independently drawn from a *U*[*0,*1] distribution. Upon division, the values of *N*_*i*_ and *D*_*i*_ are inherited by each daughter cell.

**Table 4.** Table of parameters specific to the lateral inhibition simulations.

| Parameter | Description | Model(s) | Value | Reference |
| --- | --- | --- | --- | --- |
| $t_{\text{end}}$ | Simulation duration | All | 1000 h | – |
| $p_{\text{div}}$ | Cell division rate | All | $\{0.01, 0.05, 0.1\} \text{ cell}^{-1} \text{ h}^{-1}$ | – |
| $t_{\text{min}}$ | Minimum division age | All | 1 h | – |
| $k_N$ | Dependence of Notch on Delta | All | 0.01 | [46] |
| $k_D$ | Dependence of Delta on Notch | All | 0.01 | [46] |
| $n_N$ | Notch Hill coefficient | All | 2 | [46] |
| $n_D$ | Delta Hill coefficient | All | 2 | [46] |
| $r_{DN}$ | Relative rate of Delta activity | All | $1 \text{ h}^{-1}$ | [46] |
| $R_S$ | Removal zone radius | All | 5 CD | – |
| $R_P$ | Proliferative zone radius | All | 15 CD | – |
| $r_{\text{max}}$ | Force cut-off length | OS | 1.0 | – |

Eq (14) and Eq (15) are coupled to the cell-based models using the following algorithm. At each time step, having updated the cell-based model, we calculate 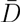 based on the current connectivity and, assuming 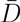 remains constant on the short interval ∆*t*, we solve the Notch signalling model numerically over the interval [*n*∆*t*, (*n* + 1)∆*t*] using a Runge-Kutta method.

Simulation snapshots for each model are shown in Fig 7. In each case, we see that lateral inhibition successfully leads to patterning of cells in ‘high delta’ steady state surrounded by cells in a ‘low delta’ steady state in the outer ring of non-proliferating cells. This patterning is disrupted in the inner proliferating region, as cells frequently change neighbours and hence are unable to synchronise their delta-notch dynamics. The degree of this disruption increases with cell division rate and is most apparent in the VM simulation. This may be due to cells exchanging neighbours more frequently, even in regions without proliferation, in the VM; a similar disruption is observed in the CP simulation. A lattice-induced anisotropy is clearly visible in the CA simulation, where cell shoving causes significantly more cell rearrangements and, as a result, less patterning along diagonals. This phenomenon also occurs, to a lesser extent, in the CP simulation.

**Fig 7.**
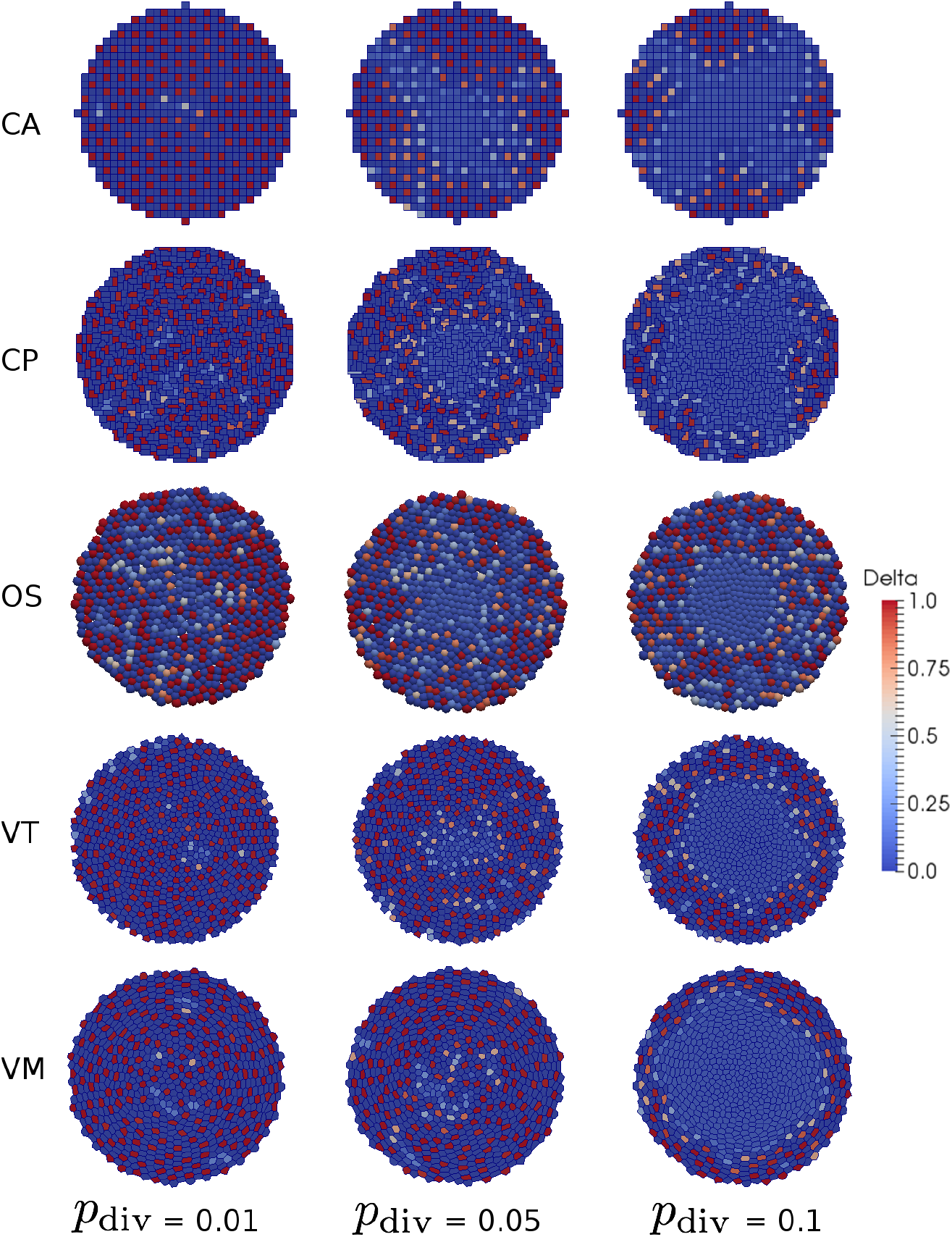
Simulations of lateral inhibition in a proliferating tissue. For each model, snapshots are shown for three levels of cell proliferation, quantified by the parameter *p*_div_. Parameter values are given in Tables 1 and 4.

A quantitative comparison of the patterning dynamics across models is shown in Fig 8. As a measure of patterning we plot the ratio of cells in the heterogeneous steady state to those not in this state at the end of each simulation, computed as a radial distribution across the tissue. Note that the ‘kinks’ observed in the CA results (Fig 8(a)) are due to the presence of discrete cells on a fixed lattice. We see that there is significantly less patterning in the proliferative region for all models and that as the rate of division is increased the difference is exaggerated.

**Fig 8.**
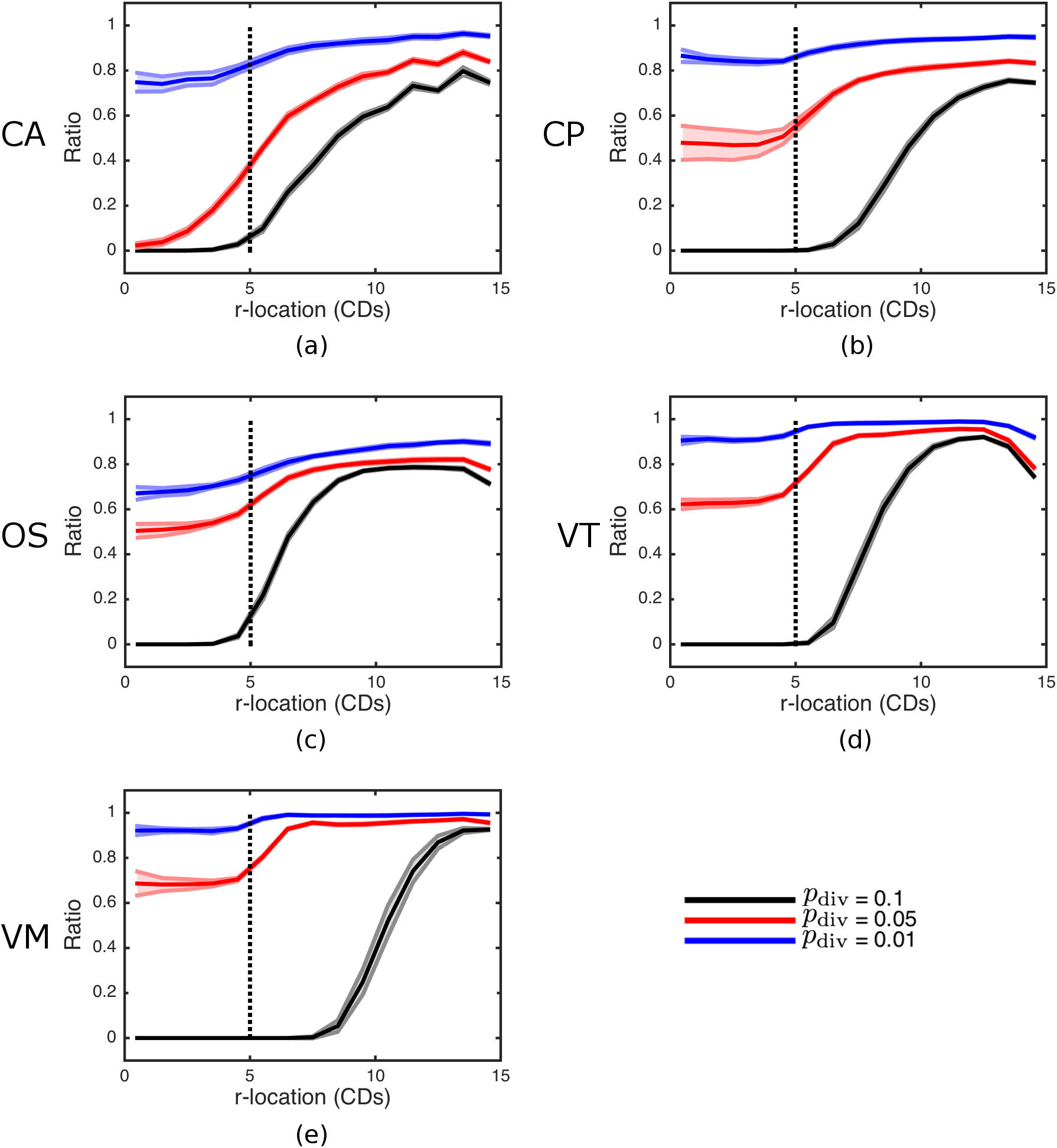
Comparison of cell fate patterning across lateral inhibition simulations. As a measure of patterning, the ratio of cells in the heterogeneous steady state to those not in this state is computed at time *t*_end_, as a radial distribution across the tissue (calculated using a bin size of 0.5 CDs), for each model: (a) CA; (b) CP; (c) OS; (d) VT; (e) VM. For each model, the mean and standard error from 10 simulations are shown for three levels of cell proliferation, quantified by the parameter *p*_div_. Parameter values are given in Tables 1 and 4.

### Long-range signalling

Morphogens are secreted signalling molecules that provide positional information to cells in a developing tissue and act as a trigger for cell growth, proliferation or differentiation. The processes of morphogen gradient formation, maintenance and interpretation are well studied, most notably in the wing imaginal disc in the fruit fly *Drosophila* [52], a monolayered epithelial tissue. A key morphogen called Decapentaplegic (Dpp) forms a morphogen gradient along the anterior-posterior axis of this tissue. Dpp is known to determine the growth and final size of the wing imaginal disc, although the mechanism by which its gradient is established remains unclear.

A number of cell-based models have been proposed for the cellular response to morphogen gradients and mechanical effects in developing tissues such as the wing imaginal disc [53, 54, 11]. As our final case study, we simulate the growth of an epithelial tissue in which cell proliferation is coupled to the level of a diffusible morphogen. This case study represents an abstraction of a wing imaginal disc and illustrates how continuum transport equations may be coupled to cell-based models.

Our description of morphogen-dependent cell proliferation is based on that proposed by [12] and is implemented as follows. The probability of a cell dividing exactly *n* time steps after its last division is given by 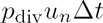, where *p*_div_ is a fixed parameter and the weighting *u*_*n*_ satisfies the recurrence relation

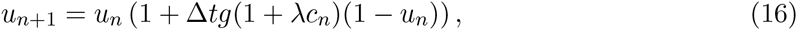

with 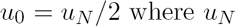 denotes the parent cell’s weighting value immediately prior to division. Here *λ* is a fixed parameter quantifying the effect of the morphogen on cell growth, *c*_*n*_ denotes the morphogen concentration at that cell at that time step, and *g* is a random variable independently drawn upon division from a truncated normal distribution with mean *µ*_*g*_, variance 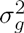 and minimum value *g*_*min*_.

When initialising the simulation, a value of *g* is drawn independently for each cell from a truncated normal distribution (as on division), and a value of *u*_0_ is drawn independently from a *U*[0.5, 1] distribution.

Each cell-based model is coupled to a continuum model of morphogen transport based on that proposed by [12]. We assume that the morphogen is secreted in a central ‘stripe’ of tissue and diffuses throughout the whole tissue, being transported by the cells, while being degraded. In this description, the morphogen concentration *c*(x, *t*) is defined continuously for times *t* ≥ 0 in the spatial domain x ∈ Ω_*t*_ defined by the boundary of the cell population (see below). This concentration evolves according to the reaction-advection-diffusion equation

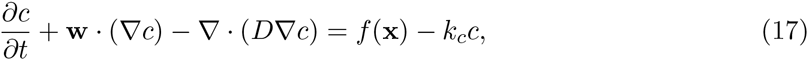

with zero-flux boundary conditions at the edge of the domain. The vector field **w** denotes the velocity of the cells moving in the tissue (and is found in the weak formulation in [12]). Its inclusion in Eq (17) denotes the advection of Dpp with the cells. The parameters D and *k*_*c*_ denote the morphogen diffusion coefficient and degradation coefficient respectively. Finally, the function *f* specifies the rate of production of morphogen in the central stripe of tissue, and is given by

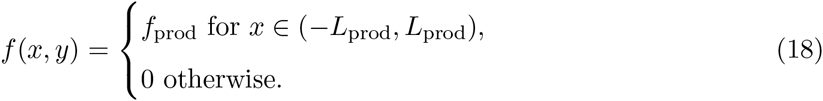

To solve Eq (17) numerically, we first discretise the spatial domain defined by the cells to make a computational mesh. For the VT model we use the triangulation defined by the dual of the Voronoi tessellation; for the vertex model we use the triangulation defined by dividing each polygon cell into a collection of triangles (made up from the set of vertices and the centre of the polygon) as in [12]; and for the CA, CP and OS models we create a triangulation by calculating the constrained Delaunay triangulation of the centres of the cells. This tessellation changes over time as the tissue grows.

We solve Eq (17) using a method of lines approach along the characteristic lines

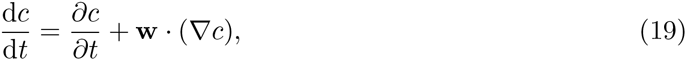

and a continuous Galerkin finite element approximation to the spatial derivatives.

We approximate the solution of Eq (17) using a Forward Euler discretization for time and a linear finite element approximation in space. As we generate the computational mesh from the cells, the mesh moves with velocity **w**. We can therefore account for the advective term of Eq (17) by moving the solution with the moving cells. Finally in each model when a cell divides it creates a new node in the mesh and the solution at the new node is defined to be the same as the node attached to the parent cell. A full list of parameter values is provided in Tables 1 and 5.

**Table 5.** Table of parameters specific to the morphogen-dependent proliferation simulations.

| Parameter | Description | Model(s) | Value | Reference |
| --- | --- | --- | --- | --- |
| $\Delta t$ | Time step | All | 0.005 h | – |
| $t_{\text{end}}$ | Simulation duration | All | 100 h | – |
| $L_r$ | Initial radius of tissue | All | 5 CD | [12] |
| $D$ | Dpp diffusion coefficient | All | $10^{-4}$ CD<sup>2</sup>h<sup>-1</sup> | [12] |
| $k_c$ | Dpp degradation rate | All | 0.01 h<sup>-1</sup> | [12] |
| $f$ | Maximal Dpp production rate | All | 0.01 h<sup>-1</sup> | [12] |
| $W_{Dpp}$ | Width of Dpp production zone | All | 4 CD | [12] |
| $p_{\text{div}}$ | Average baseline cell division rate | All | 0.1 h<sup>-1</sup> | [12] |
| $\lambda$ | Morphogen effect on cell growth | All | 0.01 | [12] |
| $f_{\text{prod}}$ | Level of production of Dpp in stripe | All | 0.01 | [12] |
| $L_{\text{prod}}$ | Half the width of stripe of Dpp production | All | 2 | [12] |
| $\mu_g$ | Cellular growth rate mean | All | 0.05 | – |
| $\sigma_g^2$ | Cellular growth rate variance | All | 0.000 1 | – |
| $g_{\text{min}}$ | Minimum cellular growth rate | All | 0.01 | – |

Simulation snapshots for each model are shown in Fig 9. As expected, over time the morphogen biases the shape of the tissue, which exhibits greater growth in the *y* direction. This is confirmed in Fig 11, which shows a quantitative comparison of tissue shape dynamics across models. A quantitative comparison of the spatio-temporal morphogen dynamics across models is shown in Fig 10. In each case, the morphogen distribution is plotted at different times as an average over the *x* direction and over 20 simulations. While the mean behaviour is conserved across models, the CA exhibits significantly greater variation about this mean. This is due to the discrete nature of cell movement, and hence morphogen advection, in these models. Looking at the snapshots in Fig 9 we see that despite being an off lattice model the VT model exhibits some regularity in shape through growth, witnessed by straighter than expected edge segments. This is due to the method for calculating connectivity in VT model and can introduce artefacts when considering freely growing domains as seen here.

**Fig 9.**
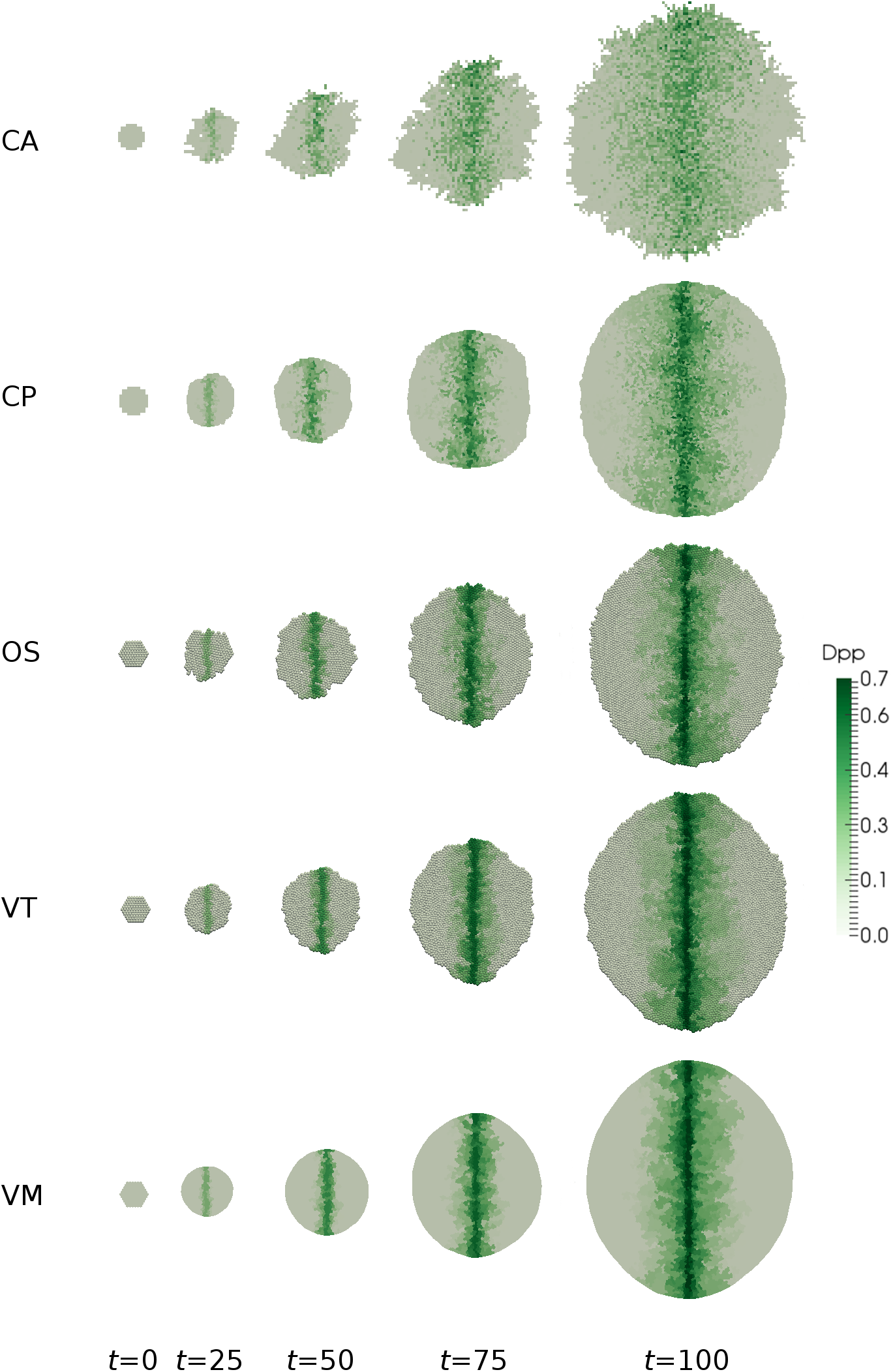
Simulations of morphogen-dependent proliferation. Snapshots of the tissue and associated morphogen distribution are shown at selected times for each model. Parameter values are given in Tables 1 and 5.

**Fig 10.**
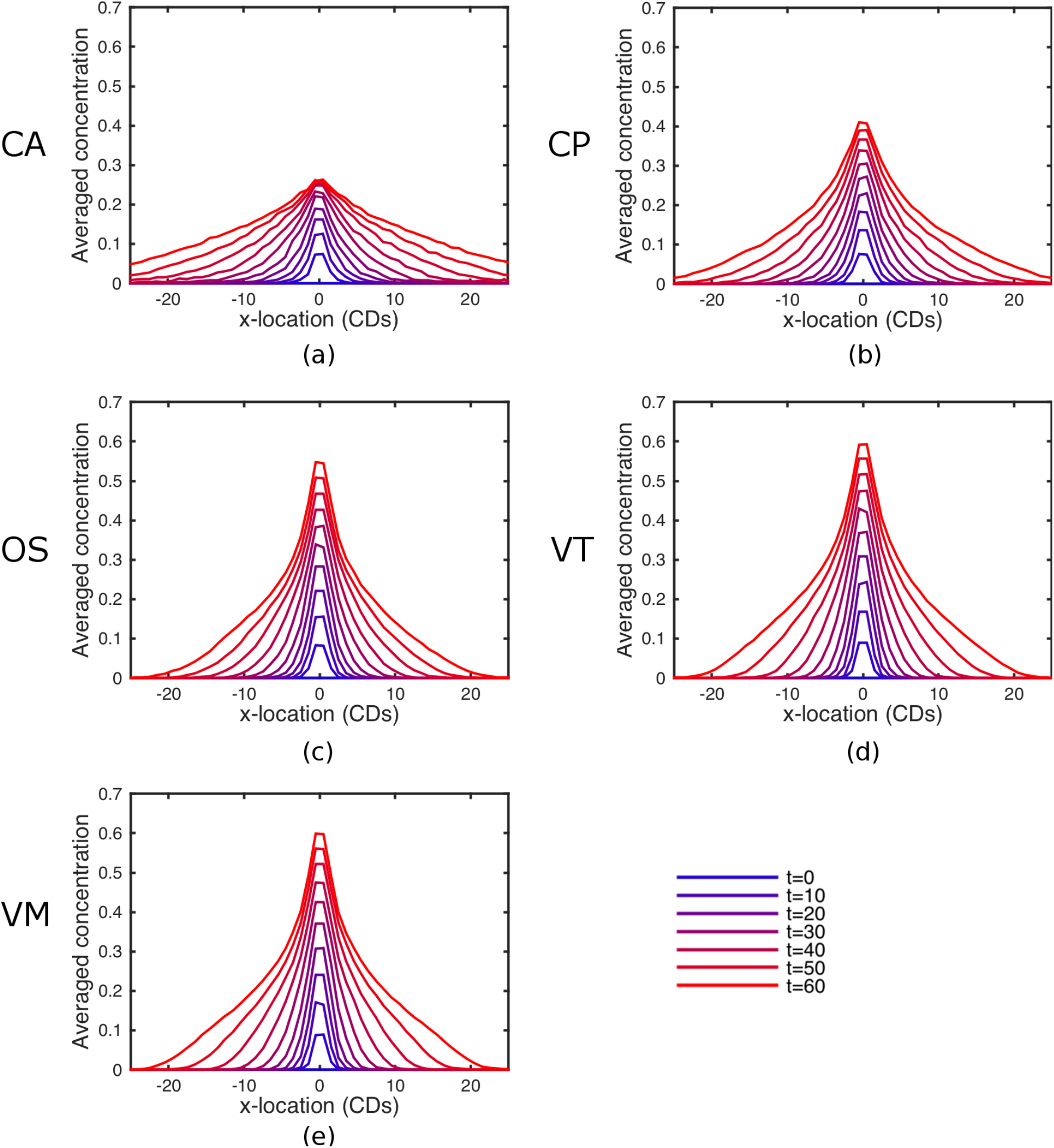
Comparison of spatio-temporal morphogen dynamics across simulations. Results are shown for each model: (a) CA; (b) CP; (c) OS; (d) VT; (e) VM. In each case, the morphogen distribution is plotted at selected times as an average over the *x* direction and over 20 simulations. Parameter values are given in Tables 1 and 5.

**Fig 11.**
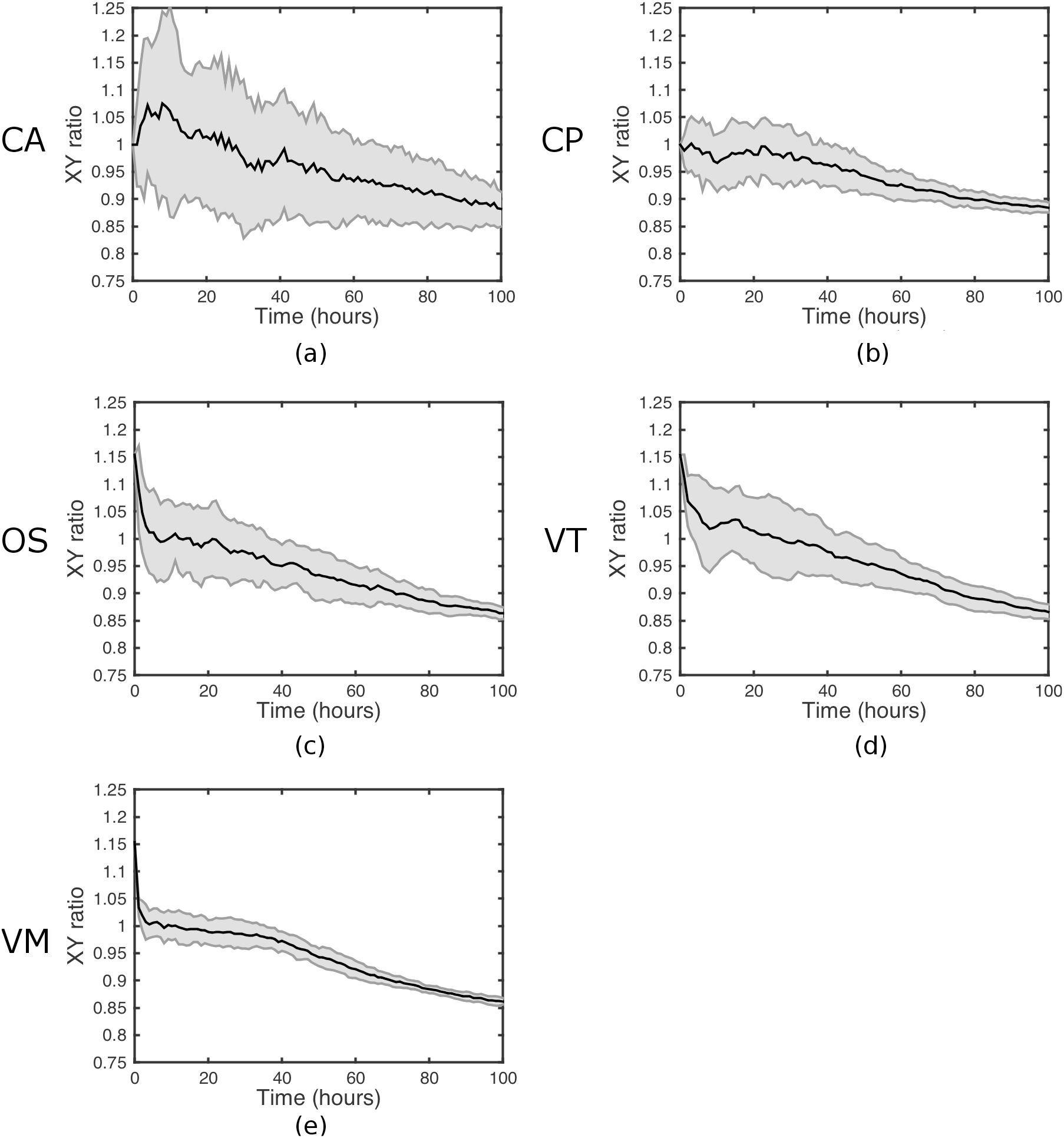
Comparison of tissue shape dynamics across morphogen-dependent proliferation simulations. As a measure of tissue anisotropy, the ratio of the widths of the tissue in the *x* and *y* directions is computed as a function of time for each model: (a) CA; (b) CP; (c) OS; (d) VT; (e) VM. For each model, we plot the mean and standard error of this ratio across 10 simulations. Parameter values are given in Tables 1 and 5.

## Discussion

The field of mathematical modelling in biology has matured beyond recognition over the past decade. One indication of this is the move towards quantitative comparison with data taking precedence over qualitative comparison. In this context, we must investigate if the model framework chosen might amplify or diminish the effects of certain processes. To this end, the present work seeks to advance our comparative understanding of different classes of models in the context of cell and tissue biology.

A variety of cell-based approaches have been developed over the last few years. These models range from lattice-based cellular automata to lattice-free models that treat cells as point-like particles or extended shapes. Such models have proven useful in gaining mechanistic insight into the coordinated behaviour of populations of cells in tissues. However, it remains difficult to accurately compare between different modelling approaches, since one cannot distinguish between differences in behaviour due to the underlying model assumptions and those due to differences in the numerical implementation. Here, we have exploited the availability of an implementation of five popular cell-based modelling approaches within a consistent computational framework, Chaste. This framework allows one to easily change constitutive assumptions within these models. In each case we have provided full details of all technical aspects of our model implementations.

We compared model implementations using four case studies, chosen to reflect the key cellular processes of proliferation, adhesion, and short- and long-range signalling. These case studies demonstrate the applicability of each model and provide a guide for model usage. While on a qualitative level each model exhibited similar behaviour, this was mainly achieved through parameter choice and fitting. Parameters were chosen to give consistent behaviour where possible. When choosing which model to use, one should bear in mind the following.

Certain examples presented in this study are more aligned with particular models. For example, in the adhesion example the CP and VM models are designed to explicitly represent cell sorting (through cell boundary energy terms) whereas the other models needed modification to represent the same phenomena. In fact, in the OS and VT (and to some extent the VM) models, the ability to sort completely was limited by the presence of local energy minima and a noise component of cell motion was required to mitigate this. However, as the level of noise was increased, artefacts can be introduced into the models, for example the tessellation may become non conformal leading to voids in the tissue.

The implementation of other features such as boundary conditions can also influence simulation outcomes. This was observed in the proliferation example where the rate of neutral drift was significantly different in the VM compared to the other models, due to additional adhesion of cells to the bottom of the domain. In this study we did not implement contact inhibition for the CA model as our definitions of contact inhibition required cells to be for different sizes. It is possible to implement an alternative form of contact inhibition in the CA model by restricting division events to only occur when there is sufficient free space [55]; however, this would again result in a different behaviour to our simulations.

A key difference between the models we considered lies in the definition of cell connectivity. It is possible for cells in the same configuration to have different neighbours under different models. For example, when under compression cells in the OS model can have more neighbours than similarly sized cells in CP, VT or VM models. The effects of this can be seen in the short-range signalling example with a high degree of proliferation.

In terms of understanding and software development time, one can code up a simple CA model in a few hours and the models increase in complexity from there in order CP, OS, VT, with VM being the most involved. While the focus here has been on epithelial layers and two spatial dimensions, all the models have also been utilised in 3D and for the CA, CP, OS, and VT, models the extension is relatively natural. For the VM model the extension is not trivial, as multiple rearrangements need to be considered in order to maintain a confluent tissue.

Finally, the models differ vastly on how long they take to simulate. In their original uncoupled forms, the least computationally complex model to simulate is the CA, followed in order by the CP, OS, VT and VM. However, this complexity depends on what is coupled to the models, at both the subcellular and tissue levels. Specifically, in order to make the CP model equivalent to the other models when coupling to subcellular and tissue level processes, we have chosen to use a time step that is smaller than that typically used in CP simulations, increasing the computation time. Table 6 shows how long typical simulations took for each example across the models considered in the present study. We see that (except for the CP model) the level of computational time is roughly as expected, increasing with complexity with the OS and VT models being similar. There are exceptions to this. For example, the CA and CP simulations of the long-range signalling example take longer than may be expected. This is due to the method used to calculate the growing PDE mesh in our computational implementation in Chaste, which is optimal for off-lattice models; future work will involve developing optimised numerical techniques that exploit the lattice structure of the on-lattice models. On the other hand, the VM simulation of the proliferation example is quicker than may be expected; this is due to the choice of parameters leading to there being slightly fewer cells in the VM simulation for this example, reducing the computational demands.

**Table 6.**
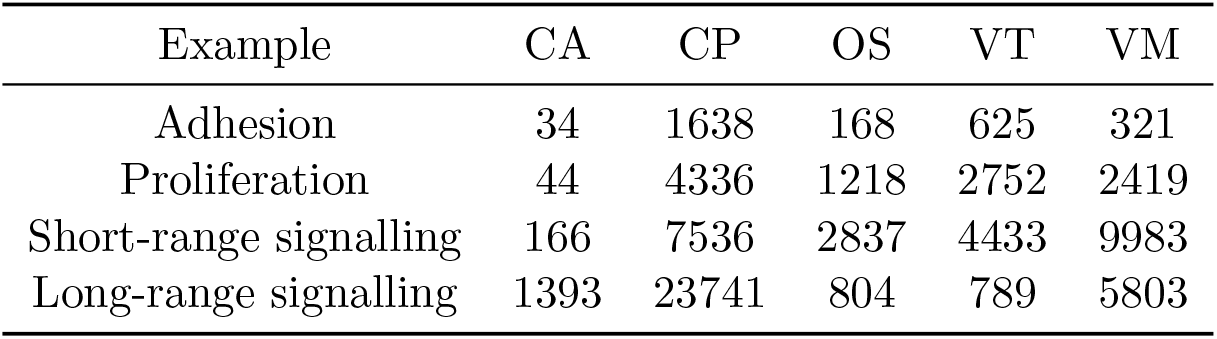
Table of approximate simulation times. Run time (in seconds, on a single core i7 processor) for typical single runs for each problem considered here.

| Example | CA | CP | OS | VT | VM |
| --- | --- | --- | --- | --- | --- |
| Adhesion | 34 | 1638 | 168 | 625 | 321 |
| Proliferation | 44 | 4336 | 1218 | 2752 | 2419 |
| Short-range signalling | 166 | 7536 | 2837 | 4433 | 9983 |
| Long-range signalling | 1393 | 23741 | 804 | 789 | 5803 |

Parallelisation is one way to both decrease computational time and to also be able to solve larger problems. Of the models considered, the CA model is simplest to parallelise. While more more advanced, the CP model has been parallelised in publicly available software packages [56], as has the OS model [57]. In the VT and VM cases, the implementations are much more involved.

The present study provides a starting point for a number of further avenues for research. First, there remains a need for theoretical and computational tools with which to easily perform quantitative model comparisons. Our results indicate that for many of the sorts of questions these types of model are currently being used to address, there is likely to be little difference in model predictions. However, such models are nevertheless moving toward a more quantitative footing, particularly as the resolution of experimental data at the cell to tissue scale improves. Further progress in this area will be accelerated by advances in automating the process of model specification and implementation, for example through extended use of mark-up languages such as SBML, FieldML and MultiCellDS.

Here we have made use of a consistent simulation framework, Chaste, within which to compare different classes of cell-based model. A longer-term challenge is to extend such comparison studies across simulation tools, of which there is an increasing ecosystem (CompuCell3d, Morpheus, EPISIM, CellSys, VirtualLeaf, Biocellion, BioFVM). We emphasize here the lack of ‘benchmarks’ on which to make such comparisons; the present study offers four examples that could offer such benchmarks.

Throughout this study we have concentrated on 2D studies. However, many of the models considered have also been implemented in three dimensions both in previous studies and in the Chaste modelling framework, for example in the case of overlapping spheres models of the intestinal crypt [33, 58] or 3D vertex models of the mouse blastocyst [59]. Of the models considered in the present study, vertex models are arguably the most technically challenging to extend to three dimensions, due to the complexity of the possible cell rearrangements and force calculations.

Work has also been done to model individual cells at a finer resolution by considering them to be composed of mesoscopic volume elements, which enables cell geometry and mechanical response to be emergent, rather than imposed, properties. These include the subcellular element model [60], which may be thought of as a natural extension of the cell centre model, and the finite element model [61] and immersed boundary model [62], which use alternative approaches to decompose cell shapes into volumetric or surface elements in a much more detailed manner than vertex models.

## Acknowledgments

JMO and AGF acknowledge funding from EPSRC through grant EP/I017909/1. The authors acknowledge the contributions made by Fergus Cooper, Jonathan Cooper, James Grogan, Daniel Harvey, Jochen Kursawe and Martin Robinson to development of the cell-based component of the Chaste software library (http://www.cs.ox.ac.uk/chaste).

